# The inattentional rhythm in audition

**DOI:** 10.1101/2024.08.10.607439

**Authors:** Troby Ka-Yan Lui, Eva Boglietti, Benedikt Zoefel

## Abstract

The detection of temporally unpredictable visual targets depends on the preceding phase of alpha oscillations (~7-12 Hz). In audition, however, such an effect seemed to be absent. Due to the transient nature of its input, the auditory system might be particularly vulnerable to information loss that occurs if relevant information coincides with the low excitability phase of the oscillation. We therefore hypothesised that effects of oscillatory phase in audition will be restored if auditory events are made task-irrelevant and information loss can be tolerated. To this end, we collected electroencephalography (EEG) data from 29 human participants (21F) while they detected pure tones at one sound frequency and ignored others. Confirming our hypothesis, we found that the neural response to task-irrelevant but not to task-relevant tones depends on the pre-stimulus phase of neural oscillations. Alpha oscillations modulated early stages of stimulus processing, whereas theta oscillations (~3-7 Hz) affected later components, possibly related to distractor inhibition. We also found evidence that alpha oscillations alternate between sound frequencies during divided attention. Together, our results suggest that the efficacy of auditory oscillations depends on the context they operate in, and demonstrate how they can be employed in a system that heavily relies on information unfolding over time.

**Significance Statement:** The phase of neural oscillations shapes visual processing, but such an effect seemed absent in the auditory system when confronted with temporally unpredictable events. We here provide evidence that oscillatory mechanisms in audition critically depend on the degree of possible information loss during the oscillation’s low excitability phase, possibly reflecting a mechanism to cope with the rapid sensory dynamics that audition is normally exposed to. We reach this conclusion by demonstrating that the processing of task-irrelevant but not task-relevant tones depends on the pre-stimulus phase of neural oscillations during selective attention. During divided attention, cycles of alpha oscillations seemed to alternate between possible acoustic targets similar to what was observed in vision, suggesting an attentional process that generalises across modalities.

## Introduction

Confronted with a dynamic environment, our brain constantly engages in the selection and prioritization of incoming sensory information. Previous research posits that neural oscillations, rhythmic fluctuations in neural excitability, are instrumental for this purpose (Schroeder & Lakatos, 2009). One fundamental assumption in this line of research is that the sensory information that coincides with the high-excitability phase of an oscillation is processed more readily than that occurring during the low-excitability phase, leading to perceptual or attentional rhythms (VanRullen, 2016b).

Previous studies in the visual modality have confirmed this assumption, demonstrating that the detection of temporally unpredictable targets depends on the pre-stimulus phase of alpha oscillations in the EEG (Busch et al., 2009; Dugué et al., 2015; Dugué et al., 2011; Mathewson et al., 2009). This phasic effect was only found for the detection of attended, but not unattended visual targets (Busch & VanRullen, 2010).

Studies in the auditory modality, however, revealed a more ambivalent role of neural oscillations in auditory perception (VanRullen et al., 2014). On the one hand, the detection of near-threshold auditory tones, presented at unpredictable moments in quiet, does not depend on pre-target neural phase (VanRullen et al., 2014; Zoefel & Heil, 2013). This result seems to question the assumption of an auditory perception that is inherently rhythmic. On the other hand, it is clear that stimulus-aligned (“entrained”) neural oscillations serve a mechanistic role in auditory attention and perception (Obleser & Kayser, 2019; Henry & Obleser, 2012; van Bree et al., 2021). Rhythmicity in auditory processing can also be observed after a cue like the onset of acoustic noise, assumed to reflect a phase reset of oscillations in the theta range (Ho et al., 2017; Lui et al., 2023; Wöstmann et al., 2020).

We here tested a hypothesis that can reconcile these apparently discrepant findings. This hypothesis is based on the fact that the auditory environment is particularly dynamic and transient (Kubovy, 1988; VanRullen et al., 2014). Losing critical auditory information that coincides with the low-excitability phase of the oscillation may be too costly for successful comprehension of auditory input. To avoid such a loss of information, the brain may therefore suppress neural oscillations in auditory system and operate in a more “continuous mode” (Schroeder & Lakatos, 2009) if incoming auditory stimuli are relevant (e.g., attended) but their timing is unknown. This assumption predicts two scenarios in which a “rhythmic mode” can be restored (Zoefel & VanRullen, 2017). First, if the timing of relevant events is known, the phase of the oscillation can be adapted accordingly, and a loss of critical information during the low-excitability phase avoided. As explained above, such an effect is fundamental for the field of “neural entrainment” (Lakatos et al., 2008, 2019). A second scenario remained unexplored and was tested here: The temporary suppression of input processing during the low-excitability phase can be tolerated if expected events are irrelevant to perform a task, even if their timing is unpredictable. In this scenario, the processing of irrelevant (but not relevant) events would be modulated by the oscillatory phase.

We measured participants’ EEG and asked them to detect pure tones at one sound frequency (task-relevant tone) and ignore pure tones at another sound frequency (task-irrelevant tone), presented at unpredictable moments (Figure 1A). We predicted that the processing of the task-irrelevant, but not that of the task-relevant tone, depends on the phase of neural oscillations. In a condition where both tones needed to be detected, we tested whether the presence of multiple task-relevant tones leads to a rhythmic alternation of attentional focus between these tones – and consequently, a phasic modulation of detection even for task-relevant tones – as previously demonstrated for the visual system (Fiebelkorn et al., 2013; Helfrich et al., 2018).

**Figure 1.**
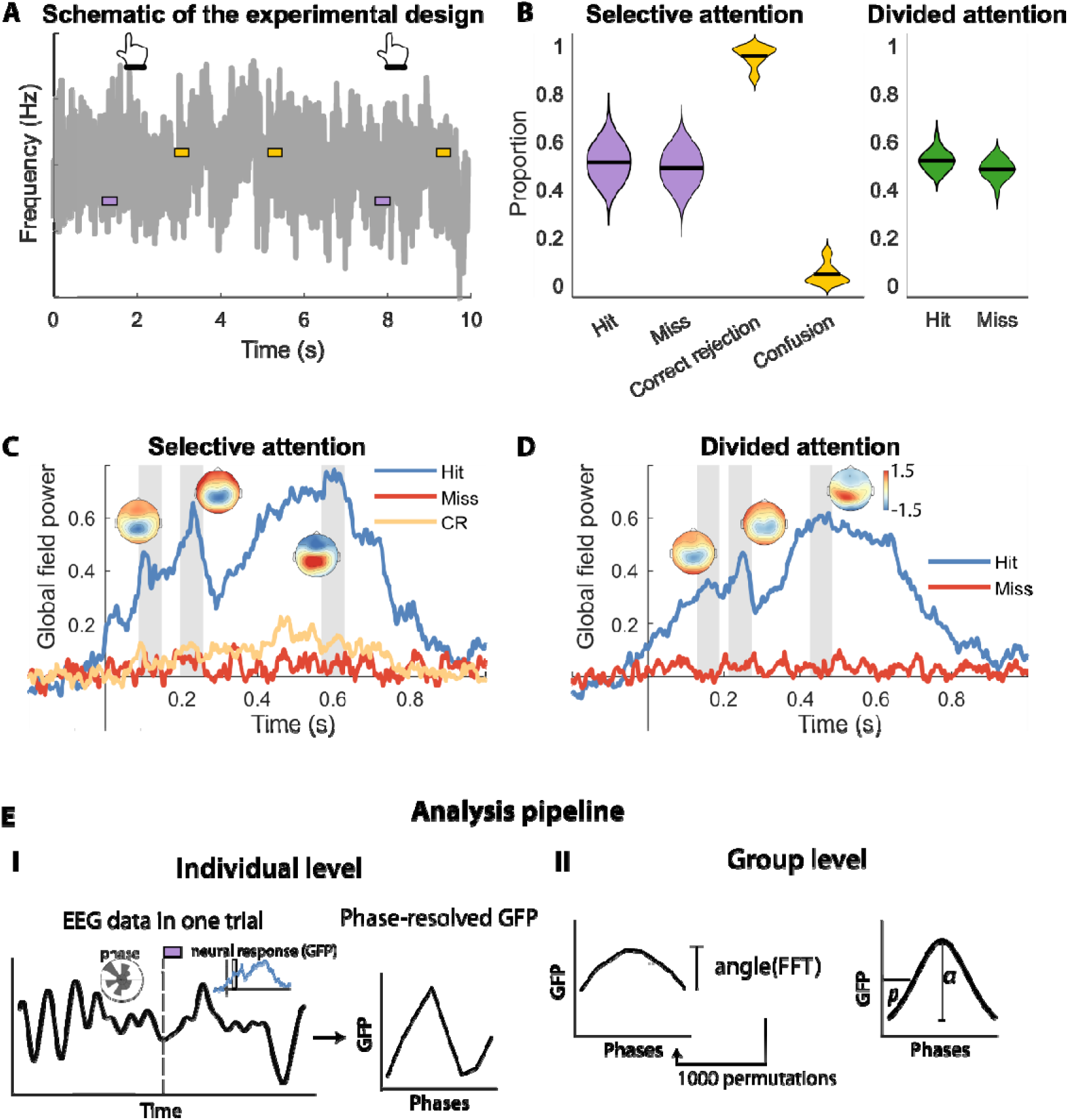
A) Schematic of the tone-in-noise detection task. Purple and yellow rectangles denote task-relevant and task-irrelevant tones, respectively. In the main experiment, low and high tones served as task-relevant and task-irrelevant tones in different blocks. Grey line shows the continuous pink noise. B) Behavioural performance for selective attention (left) and divided attention (right) conditions. Black lines show the mean across participants. C,D) Global field power (GFP) for hit (blue; relevant tones), miss (red; relevant tones), and correct rejection (CR; yellow; irrelevant tones) in the selective (C) and divided (D) attention conditions. Grey areas indicate the time window selected for the phase dependence analysis. Insets show topographies of GFP at each time window for hit trials. E) Illustration of the analysis pipeline for the phase-dependence analysis. E-I) Extraction of single-trial phase estimates for individual participants. GFP in each phase bin was calculated to create the phase-resolved GFP values. E-II) Th analysis procedure on the group level with simulated individual phase-resolved GFP for illustration (thin grey lines). The hypothesized phase effect was quantified by fitting a sine function to the averaged data (bold black line) an contrasting the amplitude *a* of this fit against that obtained in a permutation distribution (N = 1000) This analysis assumes that the phase *p* of individual sine functions is consistent across participants, an assumptions that w verified statistically (see Materials and Method; Results).

## Materials and Methods

### Participants

Thirty native French participants took part in the experiment with informed consent for a monetary reward of €25. The data of one participant was excluded due to technical issues, thus 29 participants (21 females, mean age = 22.34, SD = 1.2) were included in the final data analyses. All experimental procedures were approved by the CPP (Comité de Protection des Personnes) Ouest II Angers (protocol number 21.01.22.71950 / 2021-A00131-40).

### Experimental Design

Participants performed a tone-in-noise detection task where they were presented with pure tones at two different sound frequencies (440 Hz and 1026 Hz), embedded at unpredictable moments into a continuous stream of pink noise (Figure 1A). They were instructed to press a button when they hear a tone at the to-be-attended, task-relevant sound frequency and ignore the other one. A correct detection was defined as a button press within 1 second after pure tone onset throughout the experiment. All tones were 20ms in duration with a rise-and-fall period of 5ms. The continuous pink noise was presented at ~70 dB SPL. Prior to the main experiment, the sound level of the pure tones was titrated individually so that ~50% of tones were detected in the main task (see Adaptive Staircase Procedure). In total, 504 pure tones at each sound frequency were presented. These were divided into 12 experimental blocks, each ~ 5 min long.

In “selective attention” blocks, participants had to detect tones at one of the two sound frequencies and to ignore the other. In “divided attention” blocks, they had to detect tones at both sound frequencies. The order of the tones was pseudo-randomized with the constraint of a transitional probability between 0.24 and 0.26. The stimulus-onset asynchrony between tones was randomized between 2 and 5s with a uniform distribution to ensure temporal unpredictability. The unpredictability of the tones prevented potential preparatory responses to the upcoming stimulus. We adopted a rolling adaptive procedure to ensure that participants would detect the tone at threshold level (50%) throughout the experiment. After each block, if the participant’s detection probability was lower than 40% or higher than 60 %, the sound level of the tone at the corresponding pitch was increased or decreased by 1dB, respectively. The block order (selective attention – low pitch, selective attention – high pitch, divided attention) was counterbalanced between participants.

Stimulus presentation was done via Matlab 2019a (MathWorks, Inc., Natick, USA) and Psychtoolbox (Brainard, 1997). The auditory stimuli were presented using Etyomic ER-2 inserted earphones and a Fireface UCX soundcard. The same sound card was used to send triggers to the EEG system, ensuring synchronisation between sound and EEG.

### Adaptive Staircase Procedure

Individual detection thresholds were determined separately for each of the two pure tones with a 1-up-1-down adaptive staircase procedure as implemented in the Palamedes toolbox (Prins & Kingdom, 2018). In each adaptive trial, one pure tone was embedded randomly between 0.5 and 4.5s after the onset of a 5-second pink noise snippet. The participant had to press a button as soon as they detected the pure tone. With a starting value of −30 dB of the total soundcard output (i.e., around 70 dB SPL), the sound level of the tone decreased in steps of 1 dB if the participant correctly detected it, or increased accordingly if they missed the pure tone. The adaptive procedure ended after 10 reversals, and the final 6 reversals were used to calculate the threshold. The convergence of the staircase procedure was examined by visual inspection to determine whether the threshold would be used in the following main experiment. If convergence failed, the adaptive procedure was repeated. The average thresholds for high- and low frequency tones were −39.67 dB (SD = 1.21 dB) and −37.10 dB (SD = 1.26 dB), respectively, resulting in ~ 50% detected tones during both selective and divided attention (Figure 1B).

### EEG Recording and Data Processing

EEG was recorded using a Biosemi Active 2 amplifier (Biosemi, Amsterdam, Netherlands). 64 active electrodes positioned according to the international 10-10 system. The sampling rate of the EEG recording was 2048 Hz. Equivalent to typical reference and ground electrodes, the Biosemi system employs a “Common Mode Sense” active electrode and a “Driven Right Leg” passive electrode located in the central-parietal region for common mode rejection purposes. The signal offsets of all electrodes were kept under 50µV.

All EEG pre-processing steps were conducted using Matlab 2021a (MathWorks, Inc., Natick, USA) and the fieldtrip toolbox (Oostenveld et al., 2011). EEG data were re-referenced to the average of all electrodes. Then, the data were high- and low-pass filtered (4^th^ order Butterworth filter, cut-off frequencies 0.5 Hz and 100 Hz, respectively). Noisy EEG channels were identified by visually inspection and interpolated. Artefacts such as eye blinks, eye movements, muscle movements, and channel noise were detected in an independent component analysis (ICA) applied to down-sampled EEG data with a sampling rate of 256 Hz. Contaminated components were detected by visual inspection and removed from data at the original sampling rate. The continuous EEG data were segmented from −2s to +2s relative to each tone onset, termed “trials” in the following. Trials with an absolute amplitude that exceeded 160 µV were rejected.

We did not measure participants’ subjective perception of task-irrelevant tones as this would have rendered them relevant. Instead, we used a neural proxy to infer how readily these tones were processed, and how processing depended on pre-stimulus phase. In line with previous work (Busch & VanRullen, 2010), we used global field power (GFP) evoked by tones as such a proxy. For this purpose, event-related potentials (ERPs) were calculated for each participant, separately for correctly detected (hits) and missed targets (misses) and for each condition in the 2 x 2 design (task-relevant vs task-irrelevant, selective vs divided attention). For the selective attention condition, ERPs for trials where participants correctly did not respond to the task-irrelevant tone (correct rejection; CR) were also calculated. GFP was extracted as the low-pass filtered (cut-off frequency 10 Hz) standard deviation of the ERPs across EEG channels. Three relevant time lags for tone processing were determined as local maxima (i.e. peaks identified with the “findpeaks” Matlab function) in the grand average GFP from 0 to 1s after tone onset, separately for selective and divided conditions (Figure 1C). As the aim of this step is the identification of relevant time lags for tone processing, we restricted the analysis to detected task-relevant tones (Figure 1C, D). Time windows of interest for the analysis of phasic effects (see below) were selected as +/- 30ms around each of these three peaks. Single-trial GFP amplitudes were obtained by averaging the GFP amplitude across time points within each time window of interest. This was done separately for each experimental condition, including those without a behavioural response (i.e. the task-irrelevant conditions).

We used a fast Fourier transform (FFT) with hanning tapers and sliding windows (0.02 s steps) to extract EEG phases at frequencies from 2 Hz to 20 Hz (1 Hz steps) from single trials and channels. The window size for phase estimation was linearly spaced between 2 (for 2 Hz) and 5.6 (for 20 Hz) cycles of the corresponding frequency. The subsequent analytical steps were restricted to phases estimated from windows that do not include post-stimulus EEG data (cf. Figure 2A). This avoid a potential contamination with stimulus-evoked responses that can lead to spurious phase effects (Vinao-Carl et al., 2024; Zoefel & Heil, 2013).

**Figure 2.**
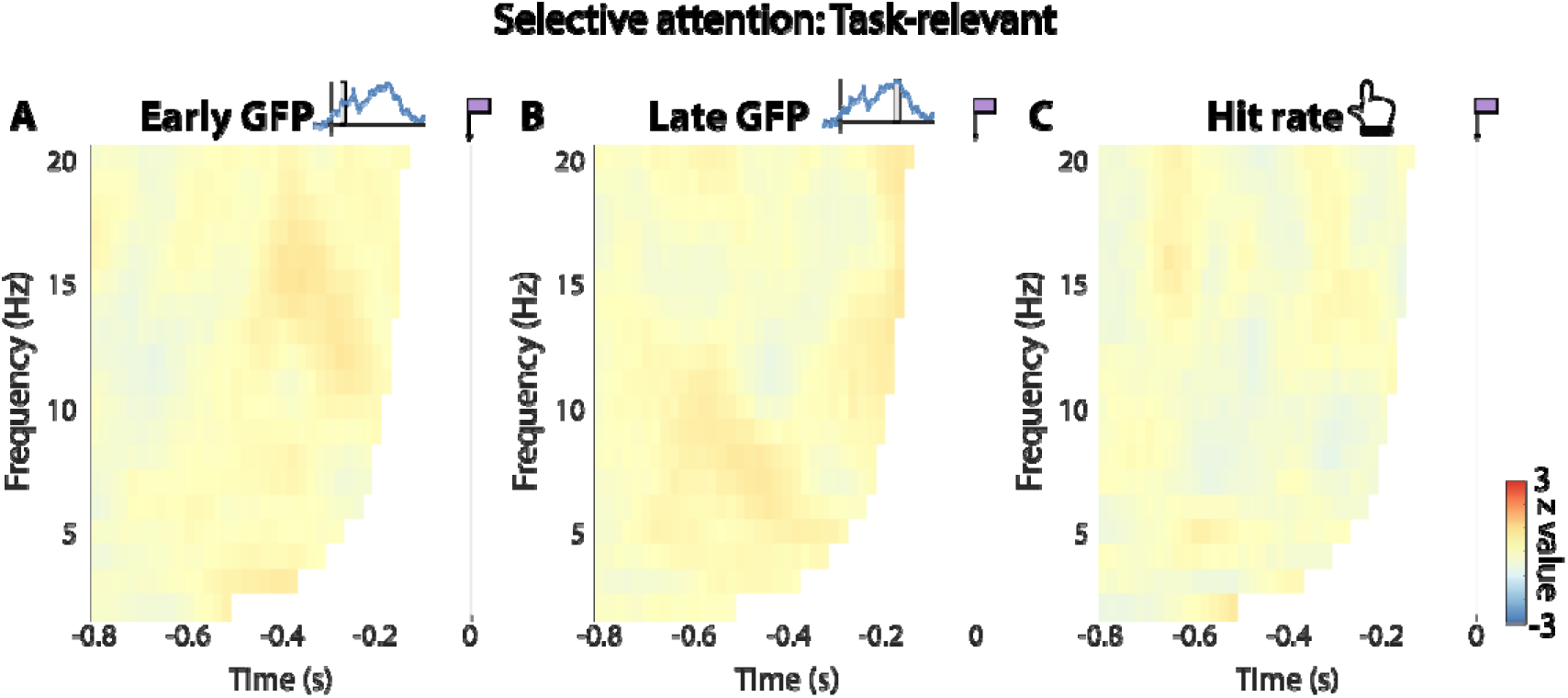
Results for task-relevant tones in the selective attention condition. The colour shows how strongly GFP (A,B) and hit rate (C) depends on EEG phase, expressed relative to a surrogate distribution, and averaged across channels. Time 0 corresponds to tone onset. In A and B, insets show relevant time lags for the analysis (early GFP: +119ms; late GFP: +598ms. Time-frequency points “contaminated” by post-stimulus data (which is “smeared” int pre-stimulus phase estimates during spectral analysis) are masked.

### Statistical analysis

To address our main hypothesis, we tested whether the magnitude of the stimulus-evoked response (as GFP; see previous section) varies with pre-stimulus neural phase (Figure 1E). We used a statistical approach that a previous simulation study (Zoefel et al., 2019) showed to be particularly sensitive to such phasic effects (“sine fit binned” method in that study). For each condition, participant, EEG channel, frequency and time point separately, single trials were divided into 8 equally spaced bins according to their phase (Figure 1E-I) and the average GFP amplitude extracted for each phase bin. We then fitted a sine function to the resulting phase-resolved GFP amplitude (Figure 1E-II). The amplitude of this sine function (*a* in Figure 1E-II) indexes how strongly tone processing is modulated by EEG phase whereas its phase (*p* in Figure 1E-II) reflects “preferred” and “non-preferred” phases for GFP (leading to highest and lowest GFP, respectively). To quantify phase effects statistically, we compared sine fit amplitudes with those obtained in a simulated null distribution, i.e. in the absence of a phasic modulation of tone processing. This null distribution was obtained by randomly assigning EEG phases to single trials and recomputing the amplitude of the sine 1000 times for each condition, EEG channel, frequency and time point (VanRullen, 2016a). For each combination of these factors, the sine amplitude from the original data was compared with the null distribution to obtain group-level z-scores:

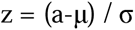

where z reflects the group level effect in the original data, a is the amplitude value in the original data, and μ and σ are mean and standard deviation (across permutations) of the subject-averaged amplitude in the surrogate distribution, respectively. Z-scores were then converted to p-values (e.g., z = 1.645 would corresponds to a significance threshold of α = 0.05, one-tailed) and corrected for multiple comparisons using the false discovery rate (FDR). Finally, clusters in combinations of frequency, time, and EEG channel for the FDR-corrected p-values were identified using the “findcluster” function in the fieldtrip toolbox.

One advantage of the statistical method used is that it makes explicit assumptions on whether participants have consistent “preferred” EEG phases, reflected in the phase of the sine fitted to individual participants (Figure 1E-II). If these phases are uniformly distributed (i.e. inconsistent across participants), the sine fit amplitude is extracted separately for each participant and then averaged before the comparison with the surrogate distribution. In this way, the z-score defined above is independent of individual preferred EEG phases. If phases are non-uniformly distributed (i.e. consistent across participants), the phase-resolved GFP (Figure 1E-I) is first averaged across participants and the sine function is fitted to the resulting average (Figure 1E-II) before the comparison with the surrogate distribution. In this way, the z-score is only high when its phase is consistent across participants. To test which version of the test is appropriate in our case, we applied a Rayleigh’s test for circular uniformity (Circular Statistics Toolbox; Berens, 2009) to the distribution of individual preferred EEG phases at each time-frequency point. We found a pre-stimulus cluster of significant phase consistency across participants (cf. Results), and adapted our statistical method accordingly (using the second version described).

We adapted this statistical approach to test whether task-relevant and task-irrelevant tones differ in their phasic modulation. In this version, we contrasted the difference in averaged sine fit amplitudes between the two conditions (relevant vs irrelevant) with another surrogate distribution for which the condition label was randomly assigned to trials. This procedure yielded another z-score which was calculated as described above.

In the divided attention condition, we additionally tested whether the processing of the low and high frequency tone has a different preferred phase by comparing the phase difference between the two tone conditions against zero (circular one-sample test against angle of 0; circ_mtest.m in the Circular Statistics Toolbox).

### Source localisation of the phase-dependence effect

We also explored the neural origins of the effects found in the analysis of EEG phase effects, using standard procedures implemented in the fieldtrip toolbox. For this purpose, we used a standard volume conduction model and electrode locations to calculate the leadfield matrix (10-mm resolution). Then, for the selective attention condition, we calculated a spatial common filter using the LCMV beamformer method (lamba = 5%; Van Veen et al., 1997) on the 20-Hz low-pass filtered EEG data from −1s to −0.5s relative to tone onset. The chosen time window encompasses all of the observed phase effects (cf. Results). This resulted in 2,015 source locations that were inside the brain.

Single-trial EEG data from individual participants were projected onto the source space with the spatial common filter. The analysis of phasic effects was then applied to data from each source location as described above for the sensor level. Due to the large computational demand, we used 100 permutations for the construction of surrogate distributions (z-score defined above), a number shown to be sufficient in the past (VanRullen, 2016a). The voxels with the 1% largest z-scores were selected as the origin of the corresponding effects on the sensory level. Note that, due to the low spatial resolution of EEG, we explicitly treat these source-level results as explorative.

## Results

### Overview

Participants were presented with tones at two different sound frequencies (Figure 1A). In some experimental blocks, they were asked to detect one of them (task-relevant tone in the selective attention condition) and ignore the other (task-irrelevant tone in the selective attention condition). In other blocks, they were asked to detect both of them (divided attention condition).

On average, participants detected 51.18% (SD = 0.07%) and 51.84% (SD = 0.04%) of task-relevant tones during selective and divided attention, respectively (Figure 1B), demonstrating successful titration of individual thresholds (see Materials and Methods).

During both attentional conditions, task-relevant tones produced a strong increase in global field power (GFP) if they were detected but not if they were missed (Figure 1C, D). We therefore used the grand-average evoked GFP as a proxy for tone processing, and identified three time lags with local GFP maxima for further analyses (grey in Figure 1C, D). The time lags for “early”, “medium” and “late” evoked GFP were 119 ms, 227 ms and 598 ms for the selective attention condition, and 159 ms, 243 ms, and 457 ms for the divided attention condition, respectively. We used GFP as a principal measure of tone processing due to the lack of behavioural response to task-irrelevant tones which would otherwise have rendered them relevant. Validating this measure of neural processing, the GFP at each of the three time lags was significantly larger for detected than for missed task-relevant tones during both selective (early: *t_28_*= 7.81, *p* < .001; medium: *t_28_* = 7.89, *p* <.001; late: *t_28_*= 10.67, *p* < .001) and divided attention (early: *t_28_* = 7.46, *p* < .001; medium: *t_28_* = 7.22, *p* <.001; late: *t_28_*= 8.44, *p* < .001).

Having identified critical time lags of tone processing, we extracted the GFP at each of the three lags evoked by single tones (including task-irrelevant ones) and tested how strongly GFP depends on pre-stimulus EEG phase in the different conditions (task-relevant vs irrelevant, selective vs divided attention). Following previous work (Lui et al., 2023; Zoefel et al., 2019), we fitted a sine function to GFP as a function of EEG phase (Figure 1E), and used the amplitude of this fit (*a* in Figure 1E-II) as a measure of phasic modulation strength. Statistical reliability of the phase effects was tested by comparison with a simulated null distribution (as z-score; see Material and Methods).

In the following, we illustrate results separately for task-relevant (Figure 2) and task-irrelevant tones (Figure 3) in the selective attention condition, respectively, as well as for the divided attention condition (Figure 4). We only display results for early and late GFP, as no phasic modulation was found for the medium time lag in any of the conditions.

**Figure 3.**
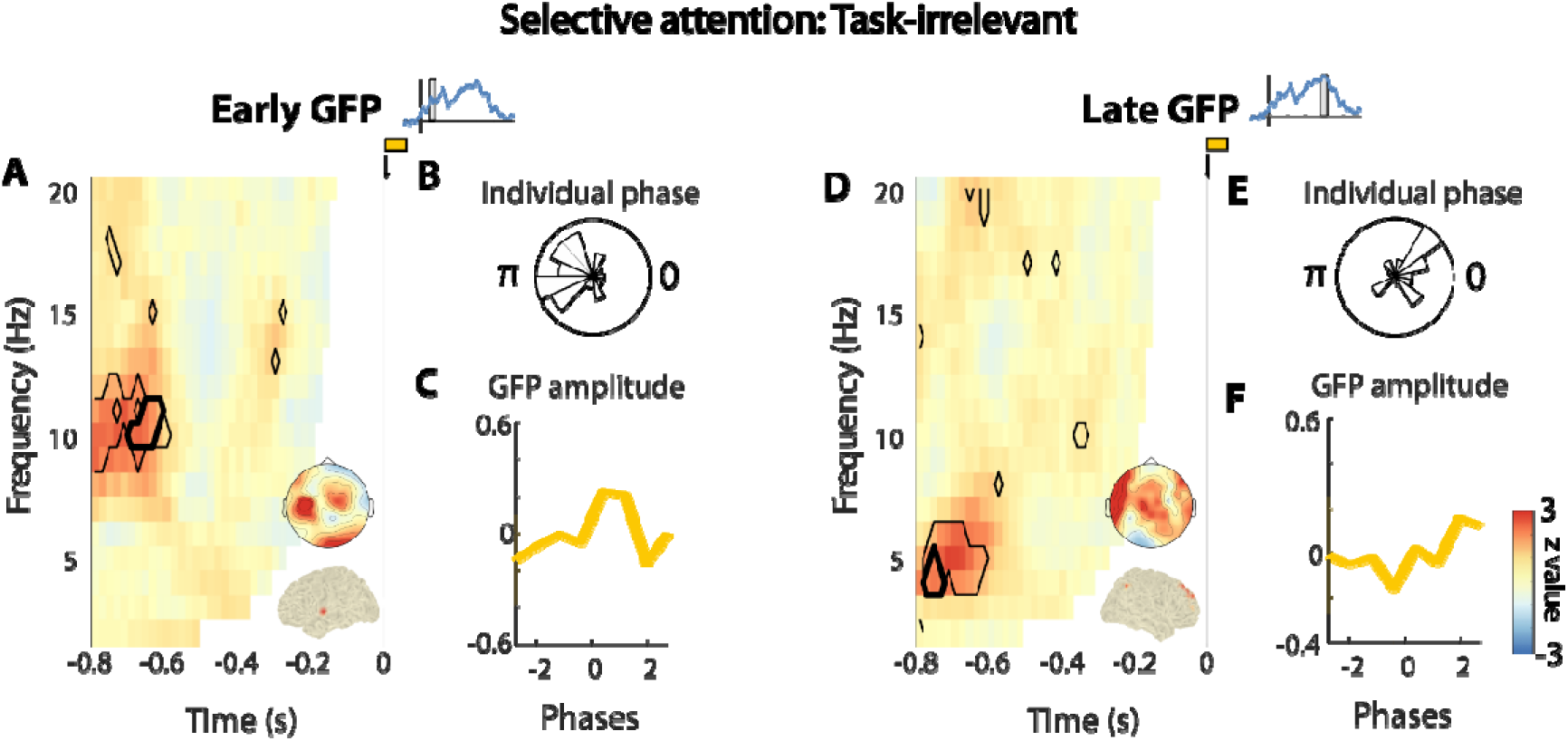
Results for task-irrelevant tones in the selective attention condition. *A, D*) Same as Figure 2A,B, but for task-irrelevant tones, and for channels selected for their significant phasic modulation of GFP (p < .05 after FDR correction). Black contours show the time-frequency points with significant phase effects. Bold black contours show the cluster with the largest summed z-score. Upper insets on the two panels show the topographical maps of z-scores in the corresponding time-frequency clusters. Lower insets show the 1% voxels with the largest source-projected z-scores in the same clusters. B, E) Distribution of individual phases of the sine function fitted to phase-resolved GFP (p in Figure 1E-II), at the time-frequency-channel combination with strongest phasic modulation (B: 11 Hz, −0.64s, C5; E: 4 Hz, −0.76s, FT7). C, F) GFP as a function of EEG phase from the same time-frequency-channel combination. The bold line shows the group-level average, the shaded area shows its standard error. Insets next t the titles show the GFP from Figure 1C with the time windows at which the analysis was performed.

**Figure 4.**
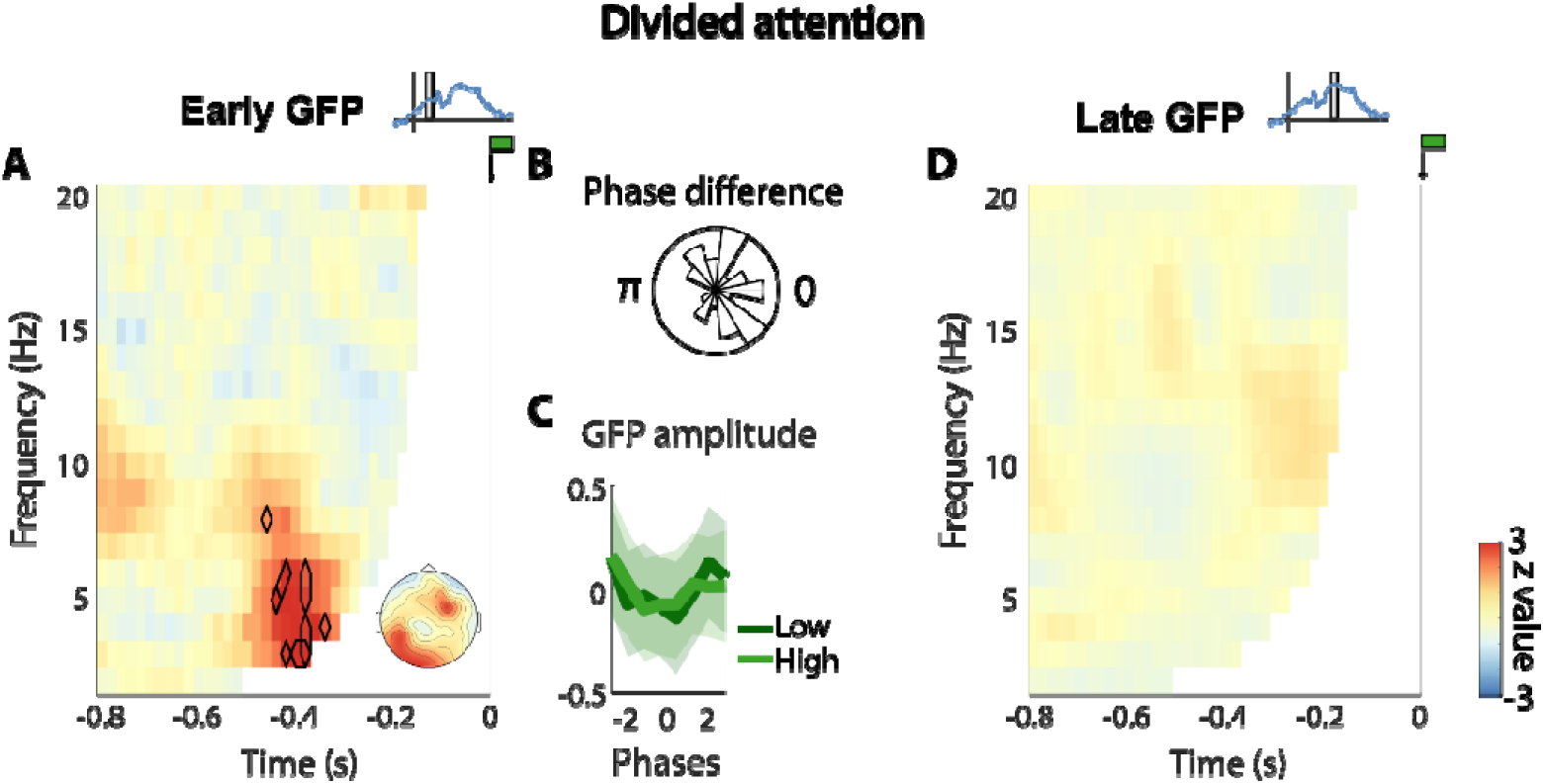
Results for task-relevant tones in the divided attention condition. All conventions as in Figure 3, apart from panel B, which illustrates the distribution of phase difference between low- and high-frequency tones, an panel C, where results are shown separately for the low- and high frequency tones.

### Neural response evoked by task-irrelevant but not task-relevant tones depends on phase of neural oscillations during selective attention

We found that pre-stimulus EEG phase did not predict GFP evoked by task-relevant tones at any of the three time lags (all *p* > 0.05 after FDR correction; Figure 2A,B). Consistent with this result, the probability of detecting these tones was independent of pre-stimulus phase (all *p* > 0.05 after FDR correction; Figure 2C). In contrast, both early (Figure 3A-C) and late (Figure 3D-F) GFP evoked by task-irrelevant tones depended on pre-stimulus phase.

For the early lag, the phasic modulation was maximal at 10 Hz and 0.8 s preceding tone onset (z = 5.41, FDR-corrected p = .003). The EEG phase leading to maximal GFP at that time-frequency point was consistent across participants (Rayleigh’s test; z = 6.21, FDR corrected *p* = .006; Figure 3B). The largest cluster of significant z-scores (FDR-corrected *p* < 0.05) was identified at ~10-11 Hz, in the left central channels, and between −0.7s and −0.62s relative to tone onset (summed z = 63.2, 14 time-frequency-channel points; Figure 3A). Explorative sourc localisation revealed that the phasic modulation originated from the left superior temporal cortex (Figure 3A, inset).

For the late lag, the phasic modulation was maximal at 5 Hz and 0.7 s preceding tone onset (z = 5.49, FDR-corrected *p* = .001). The EEG phase leading to maximal GFP at that time-frequency point was also consistent across participants (z = 6.24, FDR corrected *p* = .006; Figure 3E). The largest cluster of significant z-scores was identified at ~4-5 Hz and between −0.78 and −0.74s relative to tone onset (summed z = 47.45, 11 time-frequency-channel points; Figure 3D. This effect was localised to the right superior frontal gyrus and, to a lesser extent, the right inferior parietal cortex (Figure 3D, inset).

Contrasting amplitudes of the fitted sine functions between task-relevant and task-irrelevant tones, we found a stronger phasic modulation for the task-irrelevant tones at their relevant time-frequency points (Figure 3A,D) that concerned both early GFP (z = 4.08, *p* < .001; paired t-test) and late GFP (z = 2.92, *p* = .004). However, neither of these outcomes survived correction for multiple comparison (p > 0.05 after FDR correction).

Together, our results confirm previous findings that the processing of task-relevant auditory information is independent of the phase of neural oscillations (Zoefel & Heil, 2013), and extend them by demonstrating that such a phasic modulation reappears when the information is made irrelevant. Both alpha and theta oscillations, through their correspondence with different stages of neural processing, seem to contribute to rhythmic effects on unattended information during selective attention.

### Early but not late response evoked by task-relevant tones depends on phase of neural oscillations during divided attention

In the divided attention condition, only task-relevant tones were present. According to our principal hypothesis, the auditory system should suppress oscillations and instead operate in a continuous mode of processing to avoid a loss of information at the low-excitability phase. However, an alternative possibility is that the presence of multiple target tones requires a rhythmic alternation of attentional focus between these tones as previously demonstrated for the visual system (Fiebelkorn et al., 2013; Helfrich et al., 2018). Such a case would lead to a phasic modulation of tone processing, similarly to what we observed for task-irrelevant tones in the selective attention condition.

Figure 4 shows how strongly the evoked GFP at early (A) and late (D) time lags depended on pre-stimulus EEG phase in the divided attention condition. We found a phasic modulation of tone processing only for the early time lag. This effect was maximal at 3 Hz and 0.42 s preceding tone onset (z = 5.09, FDR-corrected *p* = .01). However, we could not identify a cluster of significant z-scores, suggesting that these did not conglomerate in neighbouring electrodes, frequency, or time as evidently as for the selective attention condition. EEG phase leading to the strongest early GFP were similar for low- and high-frequency tones (Figure 4C), supported statistically by a distribution of their phase difference (Figure 4B) that did not significantly differ from zero (mean angle = 0.23, *p* = .71; circular one-sample test against angle of 0). The probability of detecting tones did not depend on pre-stimulus phase during divided attention (all FDR corrected p < 0.05; results for time-frequency point with strongest effect in Figure 4A: z = 0.89, p = 0.37).

Together, our results show that a rhythmic mode of processing reappears in the auditory system when confronted with multiple targets, but only affects early stages of target processing. In the presence of two target tones, the frequency of modulation is approximately divided by half as compared to a single tone, and the two target tones have similar preferred EEG phases for their processing. These results speak for a mechanism processing each of the two tones at subsecutive cycles of a faster rhythm, as we explain in the Discussion.

## Discussion

The current study aimed to unveil the rhythm of auditory perception during selective and divided attention. To this end, we asked participants to perform a target-in-noise detection task where they had to attend to tones at one sound frequency and ignore another (selective attention), or had to attend to both (divided attention).

In line with previous work (Zoefel & Heil, 2013; VanRullen et al., 2014) and our own hypothesis, we found that neural and behavioural responses to task-relevant tones do not depend on the pre-stimulus phase of neural oscillations during selective attention. Conversely, early and late neural responses to task-irrelevant tones were modulated by the phase of pre-stimulus alpha and theta oscillations, respectively. These results demonstrate that while neural oscillations seem to be suppressed during attentive selection of single auditory targets, there exists a rhythmic mode of perception in the auditory system that is applied to unattended sensory information. Finally, we found evidence that this mode is also active when confronted with multiple auditory targets, although restricted to early stages of their processing.

### An inattentional rhythm in audition

It is a striking difference between modalities that selective attention *increases* the effect of neural phase on the processing of temporarily unpredictable targets in the visual domain (Busch & VanRullen, 2010) but *decreases* it in the auditory one (Zoefel & Heil, 2013; current study). Confirming previous speculations (Zoefel & VanRullen, 2017), we here demonstrate that a rhythmic mode of auditory processing is restored when stimuli become irrelevant and information loss is tolerable. This “inattentional rhythm” that seems specific to audition may arise from specific requirements on the auditory system during dynamic stimulus processing.

In contrast to the relatively stable visual environment, auditory inputs are often transient and dynamic. Therefore, periodic sampling of the external environment may be more detrimental for audition when temporarily unpredictable information is important for goal-directed behaviour. In this case, the auditory system may engage in a desynchronised cortical state in the auditory cortex that is associated with the active processing of incoming sensory inputs (Pachitariu et al., 2015). As much as this “continuous mode” prevents the loss of information by suppressing periodic moments of low excitability, it is likely to be metabolically demanding (Schroeder & Lakatos, 2009). Therefore, the auditory system may limit the use of such a mode to scenarios in which a loss of information is likely (such as the expectation of relevant events whose timing cannot be predicted). This notion can also explain the prevalence of rhythm in acoustic information (music, speech etc.): If relevant events are presented regularly, then their timing can be predicted and the oscillatory phase adapted accordingly (Lakatos et al., 2008). Such a mechanism would enable a “rhythmic mode” of processing even for task-relevant stimuli.

Based on these results, we propose that – due to its highly dynamic environment – the auditory system always needs to be “one degree more attentive” to sensory information than the visual one. We illustrate this idea in Figure 5A that can be summarised as follows: In the presence of temporarily unpredictable, relevant information, the auditory system needs to operate in a continuous mode of high-excitability, whereas the visual system can sample rhythmically, due to the significantly slower input dynamics. A similar rhythmic mode of processing is used in the auditory system to sample unattended input, whereas it is processed in a mode of continuous low sensitivity in the visual case. The latter explains why we observed a phasic modulation of task-irrelevant tones in the current study, in contrast to an absence of such an effect in the visual modality (Busch & VanRullen, 2010). Our model is also supported by the finding that auditory distractors are more distracting than visual distractors (Berti & Schröger, 2001), even when the primary task is in the visual modality (Lui & Wöstmann, 2022). This might be because the auditory system exhibits periodic moments of high sensitivity to distractors and is therefore also more sensitive to potentially threatening stimuli that warrant immediate action.

**Figure 5.**
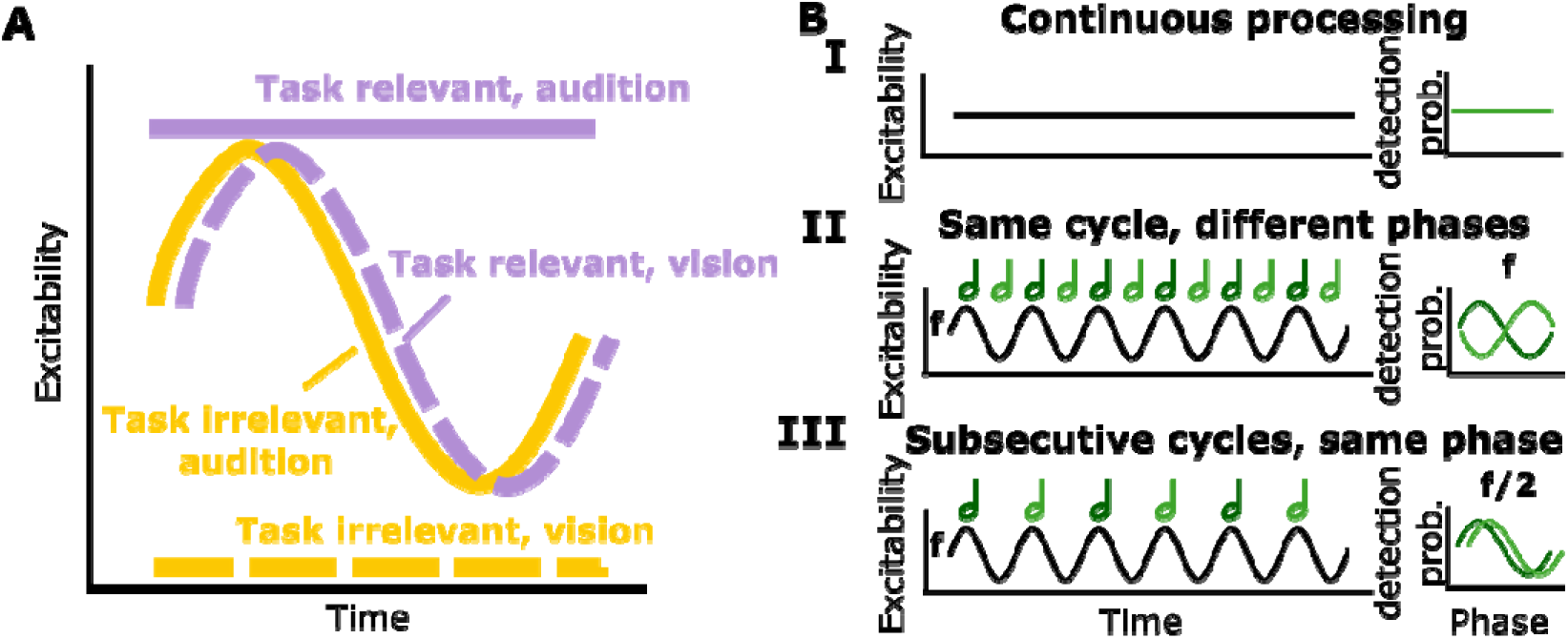
Hypothetical “modes” of processing that do or do not rely on the phase of neural oscillations durin selective (A) and divided attention (B). A) If the timing of relevant events is unknown, the auditory system might need to suppress neural oscillations to avoid a loss of information at the low-excitability phase, and operate in a mode of continuous high excitability (continuous purple line), whereas the visual system can operate rhythmically (dashed purple), due to its slower sensory dynamics. If events become irrelevant, the auditory system might chang to a mode of periodic high sensitivity, reflected in a rhythmic sampling of irrelevant information (continuous yellow). The visual system might not need these high-sensitivity moments for irrelevant information, resulting in a continuous mode of low excitability (dashed yellow). B) Three hypothetical modes of processing during auditor divided attention. When multiple targets need to be processed, the auditory system might operate in a continuous mode of processing to avoid loss of information at a low-excitability phase (I, left). Such a mode would lead to a detection of these targets that is independent of phase (I, right). Alternatively, the presence of multiple targets might require an alternation of attentional focus between possible sound frequencies that relies on neural phase at the frequency *f*. This might be achieved by prioritizing different sound frequencies at different neural phases (II, left), leading to a target detection probability that depends on the phase at *f*, and a preferred phase for detection that changes with sound frequency of the target (II, right). In an alternative rhythmic mode, possible sound frequencies are processed at the same (high-excitability) phase of f, but in subsecutive cycles (III, left), leading to a phase effect at *f/2* and to similar preferred phases across sound frequencies (III, right). The latter is what we have observed in the current study (cf. Figure 4A).

### Alpha and theta oscillations modulate distinct processing steps of irrelevant events

We found that the pre-stimulus phase of alpha oscillations predicts a relatively early response to task-irrelevant tones whereas the pre-stimulus phase of theta oscillations predicts later responses (Figure 3). We speculate that this finding can be attributed to distinct steps in the processing of task-irrelevant events that depend on different oscillatory frequency bands.

The phase of alpha oscillations is posited to gate perception via pulsed inhibition (Jensen & Mazaheri, 2010) at early stages of cortical processing where the encoding of sensory events takes place (Klimesch et al., 2011). Indeed, the phasic modulation of the early evoked response in the alpha band seemed to originate from relatively early stages of the auditory cortical hierarchy in our study (Figure 3B). The timing of the early evoked GFP (~119 ms) is well in line with components of stimulus-evoked neural responses (e.g., P1, N1) that have been associated with stimulus encoding (Näätänen & Picton, 1987). Although imaging methods with higher spatial localisation are required to validate this hypothesis, we speculate that alpha oscillations phasically modulate the encoding of task-irrelevant events (e.g., distractors).

Stimulus-evoked neural responses at later delays have been associated with higher-level cognitive operations, such as distractibility (Chao & Knight, 1995) as well as response execution and inhibition (Bokura et al., 2001). Theta oscillations in the frontal cortex have been considered a neural proxy of executive control (Mizuhara & Yamaguchi, 2007; Sauseng et al., 2007). A previous study showed evidence for a theta rhythm in distractibility by showing that perceptual sensitivity is explained by pre-distractor theta phase (Lui et al., 2023). It is thus possible that the propensity to ignore task-irrelevant events depends on pre-stimulus theta oscillations. The later timing of the theta-phase modulation in our study as well as its localisation to more frontal brain regions is in line with this assumption (Figure 3D). This effect may therefore reflect the inhibition of the processing of task-irrelevant events that occurs after their encoding. The fact that only early phasic effects were present, but the later theta-phase modulation was absent during divided attention (Figure 4) further supports this assumption, as no distractors needed to be inhibited in that condition.

It remains an open question why the strongest phase effect occurred relatively early before tone onset (~−800 to −600 ms), and earlier than what has previously been reported (Busch & VanRullen, 2010; Harris et al., 2018; Zazio et al., 2021). On the one hand, the closer to stimulus onset, the stronger is the “contamination” of phase estimates by post-stimulus data (Vinao-Carl et al., 2024; Zoefel & Heil, 2013), potentially obscuring maxima closer to tone onset. On the other hand, the earliest time points that remain unaffected by temporal smearing can be estimated precisely and do not show the strongest effects (Figures 2 – 4). Other factors might therefore play a role and need to be identified in future work. For example, it is possible that the perception and suppression of task-irrelevant auditory events is achieved through connectivity with other brain regions that eventually cascades down to the auditory system at stimulus onset.

### A rhythmic mode in auditory divided attention

We found evidence for a rhythmic mode of processing during auditory divided attention, and our results provide insights into a mechanistic implementation of such a mode. The phasic modulation of the early GFP evoked by the two tones (Figure 4A) contradicts our initial hypothesis that neural oscillations are suppressed during divided attention to task-relevant tones (Figure 5B-I). Nevertheless, the two tones (low- and high sound frequency) could be processed in the same oscillatory cycle but at different phases (Figure 5B-II) as often proposed in the context of neural oscillations (Gips et al., 2016; Jensen et al., 2014), or in subsecutive cycles (Gaillard & Ben Hamed, 2022) and at a similar phase (Figure 5B-III). Based on predicted results patterns that can distinguish these alternatives (Figure 5B, right panels), our results favour the second one, as (1) the frequency of the early modulation is divided approximately by two as compared to the processing of a single tone (compare Figure 3A and 4A); and (2) phases do not differ between the low and high frequency tones (Figure 4B,C). Therefore, our results suggest that alpha oscillations do not only modulate the processing of task-irrelevant information, but also the early stages of task-relevant processing during divided attention, alternating between possible sound frequencies of targets.

This conclusion is well in line with previous research. For instance, the frequency of visual perception decreases with increasing number of to-be-attended features (Holcombe & Chen, 2013; Schmid et al., 2022). The spotlight of attention has been posited to alternate between two locations when both are attended, dividing an overall ~8 Hz rhythm into a ~4 Hz fluctuation in perceptual sensitivity per location (Landau & Fries, 2012; Song et al., 2014; Zoefel & Sokoliuk, 2014). In the auditory modality, a similar alternation between the two ears has been reported during divided attention (Ho et al., 2017). We here extend this mechanism to an alternation between sound frequencies, supporting the previous observation that oscillatory mechanisms follow the tonotopic organisation of the auditory cortex (Lakatos et al., 2013; L’Hermite & Zoefel, 2023).

## Conclusion

By showing that the processing of task-irrelevant but not task-relevant tones depends on the pre-stimulus phase of neural oscillations during selective attention, we here provide evidence that oscillatory mechanisms in audition critically depend on the degree of possible information loss. We propose that this effect represents a crucial difference to the visual modality which might not be equally responsive to sensory information (Figure 5). During divided attention, cycles of alpha oscillations seem to alternate between possible targets similar to what was observed in vision, suggesting an attentional process that generalises across modalities.

## Acknowledgements

This study was supported by a grant from the Agence Nationale de la Recherche (ANR-21-CE37-0002). The authors thank Quentin Busson for help with data collection.

## References

Berens, P. (2009). CircStat: A MATLAB Toolbox for Circular Statistics. Journal of Statistical Software, 31, 1–21. 10.18637/jss.v031.i10

Berti, S., & Schröger, E. (2001). A comparison of auditory and visual distraction effects: Behavioral and event-related indices. Cognitive Brain Research, 10(3), 265–273. 10.1016/S0926-6410(00)00044-6

Bokura, H., Yamaguchi, S., & Kobayashi, S. (2001). Electrophysiological correlates for response inhibition in a Go/NoGo task. Clinical Neurophysiology, 112(12), 2224–2232. 10.1016/S1388-2457(01)00691-5

Brainard, D. H. (1997). The psychophysics toolbox. Spatial Vision, 10(4), 433–436. 10.1163/156856897X00357

Busch, N. A., Dubois, J., & VanRullen, R. (2009). The Phase of Ongoing EEG Oscillations Predicts Visual Perception. Journal of Neuroscience, 29(24), 7869–7876. 10.1523/JNEUROSCI.0113-09.2009

Busch, N. A., & VanRullen, R. (2010). Spontaneous EEG oscillations reveal periodic sampling of visual attention. Proceedings of the National Academy of Sciences, 107(37), 16048–16053. 10.1073/pnas.1004801107

Chao, L. L., & Knight, R. T. (1995). Human prefrontal lesions increase distractibility to irrelevant sensory inputs. Neuroreport: An International Journal for the Rapid Communication of Research in Neuroscience, 6(12), 1605–1610. 10.1097/00001756-199508000-00005

Dugué, L., Marque, P., & VanRullen, R. (2011). The Phase of Ongoing Oscillations Mediates the Causal Relation between Brain Excitation and Visual Perception. The Journal of Neuroscience, 31(33), 11889– 11893. 10.1523/JNEUROSCI.1161-11.2011

Dugué, L., Marque, P., & VanRullen, R. (2015). Theta Oscillations Modulate Attentional Search Performance Periodically. Journal of Cognitive Neuroscience, 27(5), 945–958. 10.1162/jocn_a_00755

Fiebelkorn, I. C., Saalmann, Y. B., & Kastner, S. (2013). Rhythmic sampling within and between objects despite sustained attention at a cued location. Current Biology, 23(24), 2553–2558. 10.1016/j.cub.2013.10.063

Gaillard, C., & Ben Hamed, S. (2022). The neural bases of spatial attention and perceptual rhythms. European Journal of Neuroscience, 55(11–12), 3209–3223. 10.1111/ejn.15044

Gips, B., van der Eerden, J. P. J. M., & Jensen, O. (2016). A biologically plausible mechanism for neuronal coding organized by the phase of alpha oscillations. European Journal of Neuroscience, 44(4), 2147–2161. 10.1111/ejn.13318

Harris, A. M., Dux, P. E., & Mattingley, J. B. (2018). Detecting unattended stimuli depends on the phase of prestimulus neural oscillations. The Journal of Neuroscience, 38(12), 3092–3101. 10.1523/JNEUROSCI.3006-17.2018

Helfrich, R. F., Fiebelkorn, I. C., Szczepanski, S. M., Lin, J. J., Parvizi, J., Knight, R. T., & Kastner, S. (2018). Neural mechanisms of sustained attention are rhythmic. Neuron, 99(4), 854–865.e5. 10.1016/j.neuron.2018.07.032

Henry, M. J., & Obleser, J. (2012). Frequency modulation entrains slow neural oscillations and optimizes human listening behavior. Proceedings of the National Academy of Sciences, 109(49), 20095–20100.

Ho, H. T., Leung, J., Burr, D. C., Alais, D., & Morrone, M. C. (2017). Auditory sensitivity and decision criteria oscillate at different frequencies separately for the two ears. Current Biology, 27(23), 3643–3649.e3. 10.1016/j.cub.2017.10.017

Holcombe, A. O., & Chen, W.-Y. (2013). Splitting attention reduces temporal resolution from 7 Hz for tracking one object to <3 Hz when tracking three. Journal of Vision, 13(1), 12. 10.1167/13.1.12

Jensen, O., Gips, B., Bergmann, T. O., & Bonnefond, M. (2014). Temporal coding organized by coupled alpha and gamma oscillations prioritize visual processing. Trends in Neurosciences, 37(7), 357–369. 10.1016/j.tins.2014.04.001

Jensen, O., & Mazaheri, A. (2010). Shaping Functional Architecture by Oscillatory Alpha Activity: Gating by Inhibition. Frontiers in Human Neuroscience, 4. 10.3389/fnhum.2010.00186

Klimesch, W., Fellinger, R., & Freunberger, R. (2011). Alpha Oscillations and Early Stages of Visual Encoding. Frontiers in Psychology, 2. 10.3389/fpsyg.2011.00118

Kubovy, M. (1988). Should we resist the seductiveness of the space: Time: Vision: Audition analogy? Journal of Experimental Psychology: Human Perception and Performance, 14(2), 318–320. 10.1037/0096-1523.14.2.318

Lakatos, P., Gross, J., & Thut, G. (2019). A new unifying account of the roles of neuronal entrainment. Current Biology, 29(18), R890–R905. 10.1016/j.cub.2019.07.075

Lakatos, P., Karmos, G., Mehta, A. D., Ulbert, I., & Schroeder, C. E. (2008). Entrainment of neuronal oscillations as a mechanism of attentional selection. Science, 320(5872), 110–113. 10.1126/science.1154735

Lakatos, P., Musacchia, G., O’Connel, M. N., Falchier, A. Y., Javitt, D. C., & Schroeder, C. E. (2013). The spectrotemporal filter mechanism of auditory selective attention. Neuron, 77(4), 750–761. 10.1016/j.neuron.2012.11.034

Landau, A. N., & Fries, P. (2012). Attention samples stimuli rhythmically. Current Biology, 22(11), 1000–1004. 10.1016/j.cub.2012.03.054

L’Hermite, S., & Zoefel, B. (2023). Rhythmic Entrainment Echoes in Auditory Perception. Journal of Neuroscience, 43(39), 6667–6678. 10.1523/JNEUROSCI.0051-23.2023

Lui, T. K.-Y., Obleser, J., & Wöstmann, M. (2023). Slow neural oscillations explain temporal fluctuations in distractibility. Progress in Neurobiology, 226, 102458. 10.1016/j.pneurobio.2023.102458

Lui, T. K.-Y., & Wöstmann, M. (2022). Effects of temporally regular versus irregular distractors on goal-directed cognition and behavior. Scientific Reports, 12, 10020. 10.1038/s41598-022-13211-3

Mathewson, K. E., Gratton, G., Fabiani, M., Beck, D. M., & Ro, T. (2009). To See or Not to See: Prestimulus Phase Predicts Visual Awareness. Journal of Neuroscience, 29(9), 2725–2732. 10.1523/JNEUROSCI.3963-08.2009

Mizuhara, H., & Yamaguchi, Y. (2007). Human cortical circuits for central executive function emerge by theta phase synchronization. NeuroImage, 36(1), 232–244. 10.1016/j.neuroimage.2007.02.026

Näätänen, R., & Picton, T. (1987). The N1 Wave of the Human Electric and Magnetic Response to Sound: A Review and an Analysis of the Component Structure. Psychophysiology, 24(4), 375–425. 10.1111/j.1469-8986.1987.tb00311.x

Obleser, J., & Kayser, C. (2019). Neural entrainment and attentional selection in the listening brain. Trends in Cognitive Sciences, 23(11), 913–926. 10.1016/j.tics.2019.08.004

Oostenveld, R., Fries, P., Maris, E., & Schoffelen, J.-M. (2011). FieldTrip: Open source software for advanced analysis of MEG, EEG, and invasive electrophysiological data. Computational Intelligence and Neuroscience, 2011, 156869. 10.1155/2011/156869

Pachitariu, M., Lyamzin, D. R., Sahani, M., & Lesica, N. A. (2015). State-Dependent Population Coding in Primary Auditory Cortex. The Journal of Neuroscience, 35(5), 2058–2073. 10.1523/JNEUROSCI.3318-14.2015

Prins, N., & Kingdom, F. A. A. (2018). Applying the model-comparison approach to test specific research hypotheses in psychophysical research using the palamedes toolbox. Frontiers in Psychology, 9, 1250. 10.3389/fpsyg.2018.01250

Sauseng, P., Hoppe, J., Klimesch, W., Gerloff, C., & Hummel, F. C. (2007). Dissociation of sustained attention from central executive functions: Local activity and interregional connectivity in the theta range. European Journal of Neuroscience, 25(2), 587–593. 10.1111/j.1460-9568.2006.05286.x

Schmid, R. R., Pomper, U., & Ansorge, U. (2022). Cyclic reactivation of distinct feature dimensions in human visual working memory. Acta Psychologica, 226, 103561. 10.1016/j.actpsy.2022.103561

Schroeder, C. E., & Lakatos, P. (2009). Low-frequency neuronal oscillations as instruments of sensory selection. Trends in Neurosciences, 32(1), 9–18. 10.1016/j.tins.2008.09.012

Song, K., Meng, M., Chen, L., Zhou, K., & Luo, H. (2014). Behavioral Oscillations in Attention: Rhythmic α Pulses Mediated through θ Band. Journal of Neuroscience, 34(14), 4837–4844. 10.1523/JNEUROSCI.4856-13.2014

van Bree, S., Sohoglu, E., Davis, M. H., & Zoefel, B. (2021). Sustained neural rhythms reveal endogenous oscillations supporting speech perception. PLOS Biology, 19(2), e3001142. 10.1371/journal.pbio.3001142

Van Veen, B. D., Van Drongelen, W., Yuchtman, M., & Suzuki, A. (1997). Localization of brain electrical activity via linearly constrained minimum variance spatial filtering. IEEE Transactions on Biomedical Engineering, 44(9), 867–880. 10.1109/10.623056

VanRullen, R. (2016a). How to Evaluate Phase Differences between Trial Groups in Ongoing Electrophysiological Signals. Frontiers in Neuroscience, 10. 10.3389/fnins.2016.00426

VanRullen, R. (2016b). Perceptual Cycles. Trends in Cognitive Sciences, 20(10), 723–735. 10.1016/j.tics.2016.07.006

VanRullen, R., Zoefel, B., & Ilhan, B. (2014). On the cyclic nature of perception in vision versus audition. Philosophical Transactions of the Royal Society B: Biological Sciences, 369(1641), 20130214. 10.1098/rstb.2013.0214

Vinao-Carl, M., Gal-Shohet, Y., Rhodes, E., Li, J., Hampshire, A., Sharp, D., & Grossman, N. (2024). Just a phase? Causal probing reveals spurious phasic dependence of sustained attention. NeuroImage, 285, 120477. 10.1016/j.neuroimage.2023.120477

Wöstmann, M., Lui, T. K.-Y., Friese, K. H., Kreitewolf, J., Naujokat, M., & Obleser, J. (2020). The vulnerability of working memory to distraction is rhythmic. Neuropsychologia, 146, 107505. 10.1016/j.neuropsychologia.2020.107505

Zazio, A., Ruhnau, P., Weisz, N., & Wutz, A. (2021). Pre-stimulus alpha-band power and phase fluctuations originate from different neural sources and exert distinct impact on stimulus-evoked responses. European Journal of Neuroscience, ejn.15138. 10.1111/ejn.15138

Zoefel, B., Davis, M. H., Valente, G., & Riecke, L. (2019). How to test for phasic modulation of neural and behavioural responses. NeuroImage, 202, 116175. 10.1016/j.neuroimage.2019.116175

Zoefel, B., & Heil, P. (2013). Detection of near-threshold sounds is independent of EEG phase in common frequency bands. Frontiers in Psychology, 4. 10.3389/fpsyg.2013.00262

Zoefel, B., & Sokoliuk, R. (2014). Investigating the Rhythm of Attention on a Fine-Grained Scale: Evidence from Reaction Times. Journal of Neuroscience, 34(38), 12619–12621. 10.1523/JNEUROSCI.2134-14.2014

Zoefel, B., & VanRullen, R. (2017). Oscillatory mechanisms of stimulus processing and selection in the visual and auditory systems: State-of-the-art, speculations and suggestions. Frontiers in Neuroscience, 11, 13.

